# Neural Circuit Mechanism of Decision Uncertainty and Change-of-Mind

**DOI:** 10.1101/377432

**Authors:** Nadim A A Atiya, Iñaki Rañó, Girijesh Prasad, KongFatt Wong-Lin

## Abstract

Decision-making is often accompanied by a degree of confidence on whether a choice is correct. Decision uncertainty, or lack in confidence, may lead to change-of-mind. Studies have identified the behavioural characteristics associated with decision confidence or change-of-mind, and their neural correlates. Although several theoretical accounts have been proposed, there is no neural model that can compute decision uncertainty and explain its effects on change-of-mind. We propose a neuronal circuit model that computes decision uncertainty while accounting for a variety of behavioural and neural data of decision confidence and change-of-mind, including testable model predictions. Our theoretical analysis suggests that change-of-mind occurs due to the presence of a transient uncertainty-induced choice-neutral stable steady state and noisy fluctuation within the neuronal network. Our distributed network model indicates that the neural basis of change-of-mind is more distinctively identified in motor-based neurons. Overall, our model provides a framework that unifies decision confidence and change-of-mind.

---

The decisions we make are often accompanied with a degree of uncertainty – how likely a decision will be correct^1–3^. Some decisions are more difficult than others, inducing internal conflict that may lead to reconsideration or change-of-mind^4,5^. Likewise, challenging decisions are associated with higher uncertainty, more errors and longer response times^1,6,7^. This high uncertainty could also result in subsequent behavioural adjustments, affecting how quickly and accurately we make consecutive decisions^8,9^. Several theoretical and experimental accounts posit that uncertainty is computed while making decisions^6,7,10–15^. However, how decision uncertainty is encoded in the brain and the neural mechanism by which it affects changes-of-mind and subsequent behavioural adjustments has, so far, remained elusive^16–18^.

The neural correlates of decision uncertainty have been gradually revealed in animal and human studies^6,7,13,19–21^. For instance, neural recordings from animals demonstrated strong correlation between lower-rate neuronal firing activity in the lateral intraparietal area (LIP) of the cortex and high decision uncertainty^7^. Computational models have accounted for this, suggesting that neural responses are represented by probability distributions, where uncertainty can be quantified by evaluating the posterior probability^10,22^. These models, however, imply Bayesian optimality^23^, with no consensus on how this optimality emerges from the neurobiology^8,24^.

Other experimental studies have shown weaker linkage between choice accuracy and uncertainty-level reporting^6,11,19,25,26^. For instance, patients with lesions in the prefrontal cortex demonstrated poor confidence reporting performance, while choice accuracy was largely unaffected^19^. Several computational models support this view by predicting a dissociation between uncertainty and the formation of a perceptual decision^27,28^. For instance, in one model^27^, an extension of the drift-diffusion decision-making model (for evidence accumulation) ^29,30^, the evidence accumulation continues after a decision is reached, and hence a post-decision confidence rating can be provided. Specifically, the parameters controlling the post-decision stage are independent from the ones that control initial decision processing stage.

Changing one’s mind has been attributed to processing new evidence that negates a previous judgement^4^. More recent neurophysiological evidence has shown that some changes-of-mind occur as a result of an internal error-correction mechanism^25^, suggesting decision uncertainty plays a role in inducing changes-of-mind^31^. However, the neural mechanism of decision uncertainty (within a single trial or across consecutive ones) and its link to change-of-mind has so far remained ambiguous. In particular, there is no neural circuit model that explains this shared neural mechanism^17^.

Within the studies of perceptual decision confidence/uncertainty and change-of-mind, there are some common findings that have been identified (Supplementary Figs. 1 and 2). Firstly, more difficult tasks, associated with lower (sensory) evidence quality, lead to higher decision uncertainty, which is also associated with lower choice accuracy (Supplementary Fig. 1) ^6,32^. Secondly, higher decision uncertainty is associated with lower evidence quality for correct choices while counter-intuitively associated with better evidence quality for incorrect choices (forming the often observed “<” pattern) (Supplementary Fig.1) ^6,11,33,34^. Thirdly, changes-of-mind are more likely to occur when the task is more difficult, and more often accompanied by correcting an initial impending error choice – hence more error-to-correct changes than correct-to-error changes^4,35^ (although the difference has been shown to vary in some cases^35^). Further, the likelihood of correct changes-of-mind (to the subsequent correct choices) may peak at an intermediate level of task difficulty and then decrease gradually when the task becomes much easier (Supplementary Fig. 2) ^4,35^.

**Figure 1.**
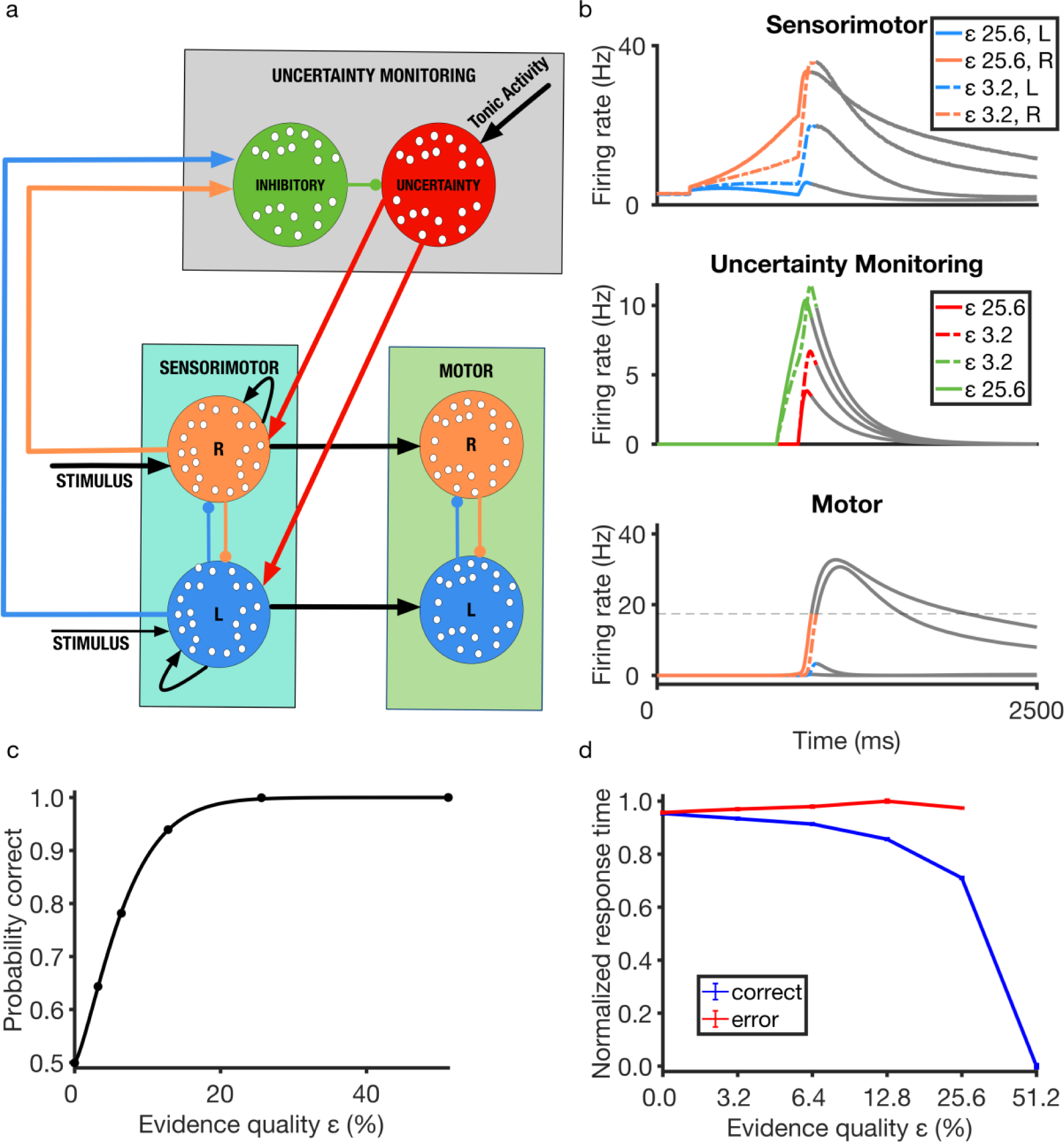
Schematic diagram and performance of the distributed neural circuit model. (**a**) The model consists of three modules. The uncertainty-monitoring module consists of two neuronal populations. Inhibitory neuronal population (green) receives excitatory input (straight arrows) from output of sensorimotor module while inhibiting the uncertainty-encoding neuronal population (lines with filled circles), which in turn provides excitatory feedback to sensorimotor module. The uncertainty-encoding population receives a constant tonic excitatory input which varies across trials in specific cases (i.e. multi-stage paradigm, see Methods and below). The sensorimotor module consists of two competing (mutually inhibitory) neuronal populations each selective to noisy sensory information (e.g. rightward or leftward random-dot motion stimulus) favouring one of two (e.g. right R or left L) choice options. The motor module, receiving inputs from sensorimotor module, also consist of neural integrators that report the choice made (**b**) Timecourse of neuronal population firing rates averaged over non-change-of-mind trials with evidence quality, *ε* = 25.6% (easy task; solid lines) and *ε* = 3.2% (difficult task; dashed lines), where *ε* is equivalent to motion coherence in the classic random-dot stimulus. Faster ramping activity (top and bottom panels) with lower uncertainty quantification (middle panel; red) with larger *ε*. Colour of activity traces reflects the associated neural populations in (a). To reveal the full network dynamics, the network activities (greyed out) were not reset after a choice was made. (**c**) Psychometric function used to fit choice accuracy (using a Weibull function, see Methods). (**d**) Response times for correct (blue) and error (red) responses from the model. In this example, the activation onset times for the inhibitory and uncertainty-encoding neuronal populations are 400 ms and 500 ms after stimulus onset, respectively.

**Figure 2.**
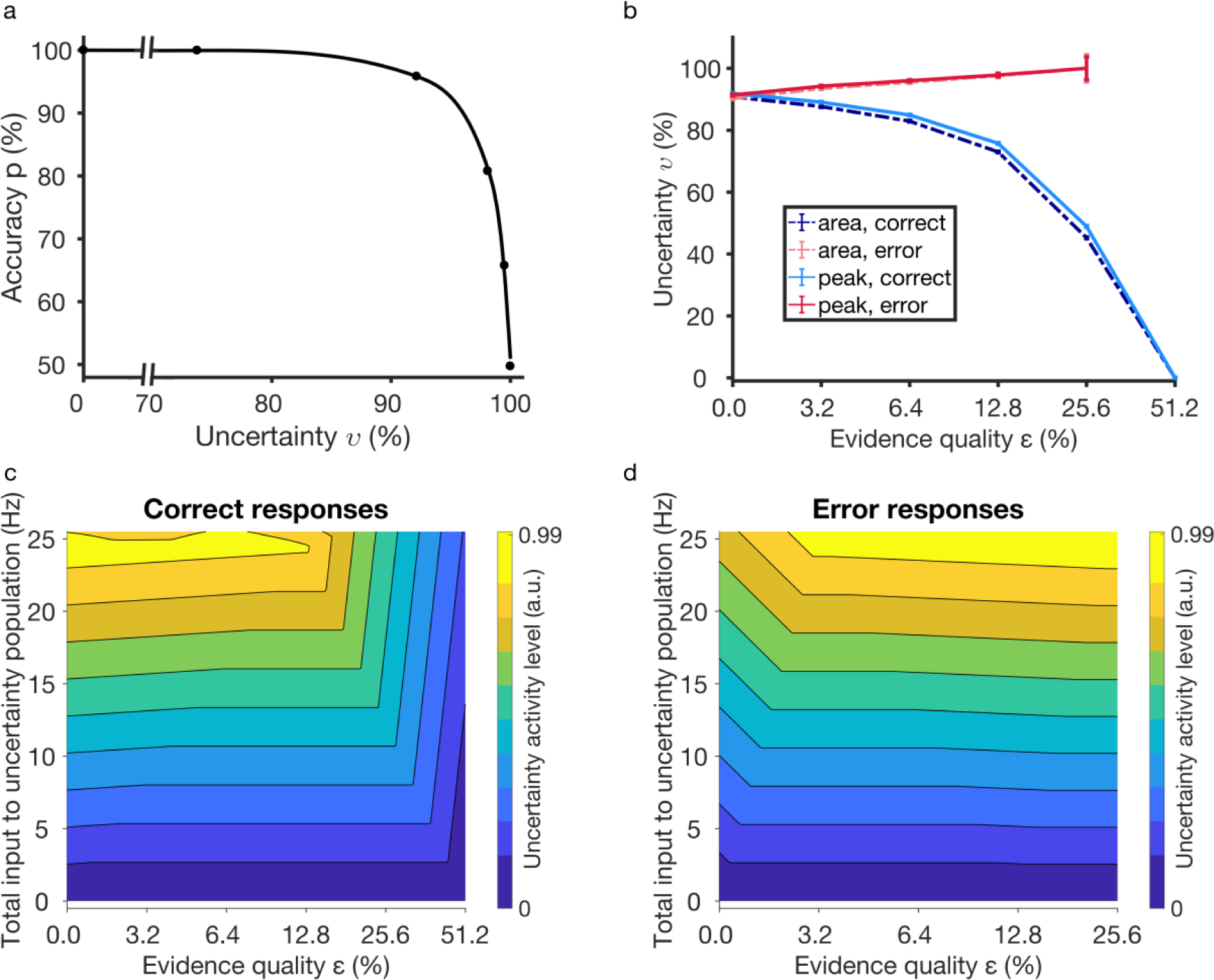
Model accounts for relationships among decision uncertainty, accuracy and evidence quality. (**a**) Choice accuracy as a function of decision uncertainty (based on peak value of uncertainty-encoding neuronal population activity). (**b**) Decision uncertainty as a function of evidence quality *ε*. Red (blue): error (correct) choices. Bold (dashed): Uncertainty measure based on averaged peak (peak) or temporal integral (area) of the uncertainty-encoding neuronal population activity (Methods). Error bars are SEM. (**c**-**d**) Activity level of uncertainty-encoding population depends on the total input to the uncertainty-encoding population and evidence quality. Uncertainty activity level is normalised (see Methods). (c) Correct responses. Activity of uncertainty-encoding population is higher *ε*) due to prolonged response times (RTs) (Fig. 1d), allowing the uncertainty-encoding population longer time to integrate. See text for more detailed description. (d) Error responses. Activity of uncertainty-encoding population is higher during errors in easier tasks (higher *ε*) due to prolonged RTs (Fig. 1d), allowing the uncertainty-encoding population longer time to integrate.

In this work, and to the best of our knowledge, guided by the above findings and related neural data (Supplementary Fig. 3), we have developed the first cortical neural circuit computational model that can mechanistically quantify and monitor decision uncertainty, which may subsequently cause a change-of-mind, hence unifying the two areas of study. Our multi-layer recurrent network model not only accounts for the abovementioned key characteristics of decision uncertainty^6,10,36^ and change-of-mind^4,35^ across a wide variety of experiments (of both behavioural and neural data) but also sheds light on their neural circuit mechanisms. In particular, using dynamical systems analysis, we show that change-of-mind occurs due to the presence of a transient choice-neutral stable steady state together with noisy fluctuations within the neuronal network. Interestingly, because our model consists of multiple layers of neural integrators, we found that the reversal of competing neural activity encoding the choices (neural basis for change-of-mind) is more likely to be more distinctive for neurons near the motor execution area, without necessarily requiring a clear reversal of neural activity at more upstream sensory or sensorimotor neurons.

**Figure 3.**
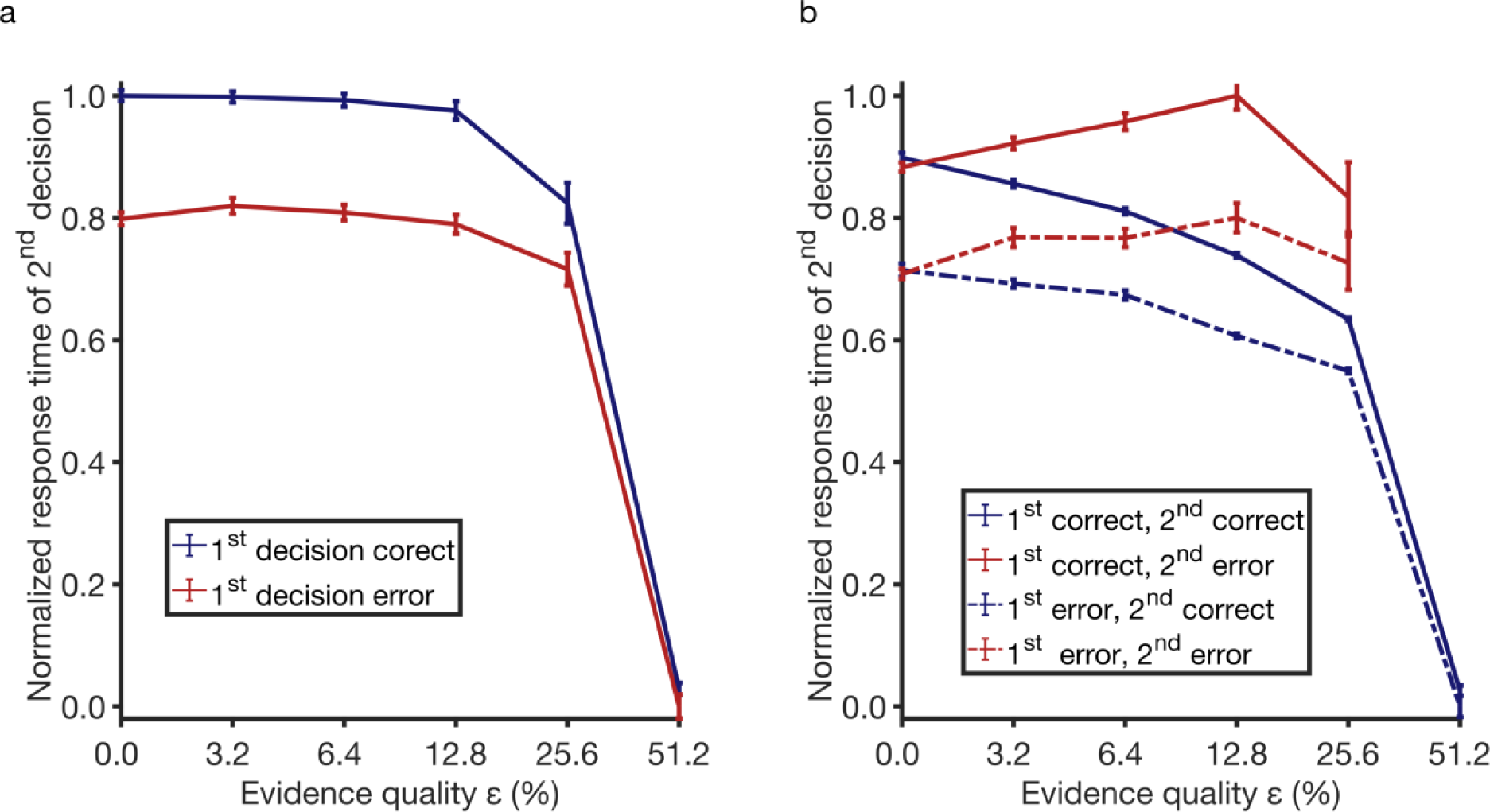
Model accounts for and predicts how response times are affected by uncertainty in the previous decision. (**a**) Normalized response times of the 2^nd^ decision when the first decision is correct (blue) and error (red). The model exhibits faster response times, when the 1^st^ decision is error compared to correct replicating experimental observations^43^. (**b**) Response times of the second decisions are further split into error (red) and correct (blue) decisions. Model predicts a slightly bigger difference between 2nd error responses (bold vs. dashed red lines) than 2nd correct responses (bold vs. dashed blue lines). Error bars are SEM.

## Results

### Neural circuit model computes decision uncertainty

We propose a novel neural circuit model that can encode, quantify, and monitor decision uncertainty, which we named the decision uncertainty-monitoring module (Fig. 1a, grey box). This circuit is built on our previous biologically-motivated neural circuit model of decision-making that focuses on sensory evidence accumulation^37^ (Fig. 1a).

The uncertainty-monitoring module receives input based on the summed sensorimotor neuronal populations activities (Fig. 1a and b). In particular, a population of inhibitory neurons (Fig. 1a, green circle) integrates these summed activities (Fig. 1a, blue and orange pointed arrows; Methods). This neuronal population in turn inhibits a neighbouring excitatory neuronal population that encodes decision uncertainty (Fig. 1a, red circle). Hence, decision uncertainty can be continuously monitored (Fig. 1b, middle). Together, the network structure with these two neuronal populations is reminiscent of a cortical column^38^.

Further, decision uncertainty information from the uncertainty-monitoring module is continuously fed back equally to the sensorimotor neuronal populations (Fig. 1a, light blue box), thus providing, effectively, an excitatory feedback mechanism between the two brain systems, which consequently may affect the final decision outcome, and in some instances, even lead to change-of-mind, as we shall demonstrate below. This feedback loop, as in control theory, provides the key computational basis of linking decision uncertainty and change-of-mind. Without this feedback loop, the model does not exhibit change-of-mind behaviour (Supplementary Fig. 4). However, it can still encode decision uncertainty and produce the experimentally-observed relationship between decision uncertainty and task difficulty (Supplementary Fig. 5). In addition, the neural circuit model also has motor-based neuronal populations either located within the same brain region or downstream in the decision-processing pathway (Fig. 1a, green box). Inputs to these populations are temporally integrated based on the neural firing rate outputs of the associated sensorimotor neuronal populations (Fig. 1b, bottom; Methods).

**Figure 4.**
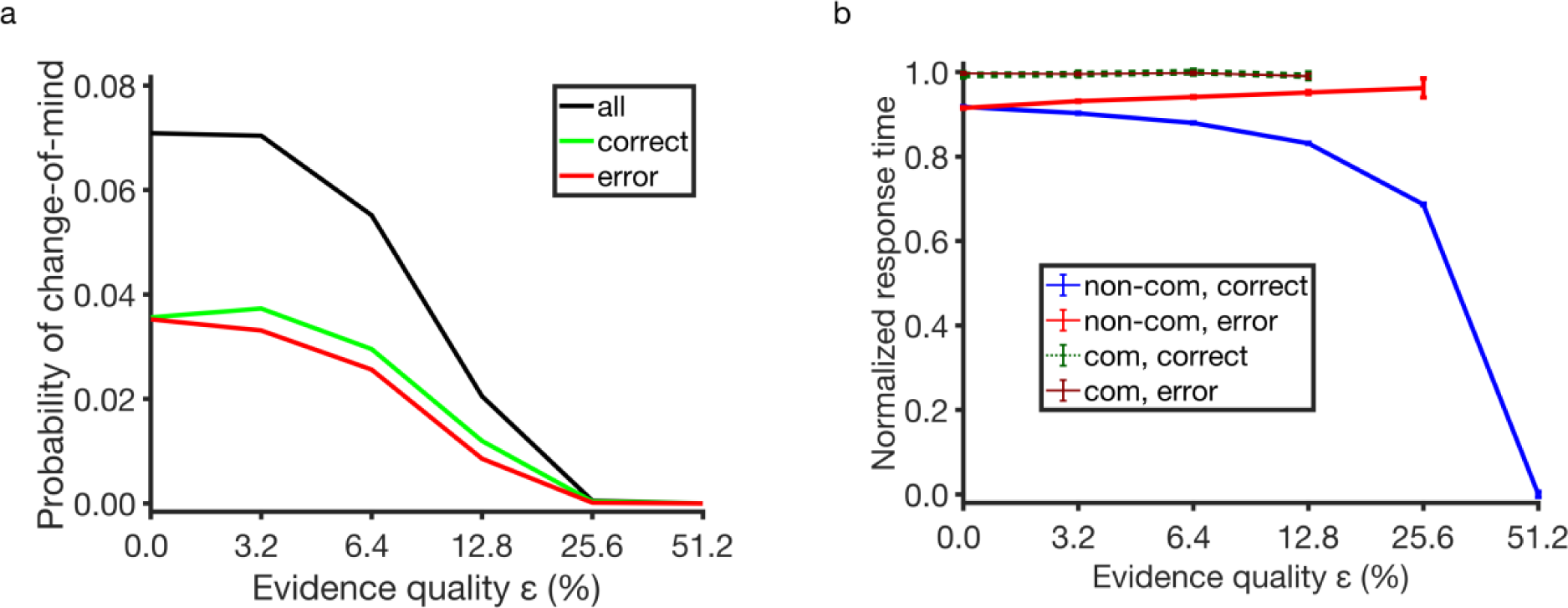
Model accounts for and predicts key characteristics of change-of-mind. (**a**) Probability of change-of-mind with respect to evidence quality. Probability of change-of-mind for a single evidence quality level is calculated by dividing the total number of change-of-mind trials by the total number of simulated trials for a specific evidence quality (see Methods). Black: Total probability of change-of-mind, consisting of both correct and error choices. Green (red): only subsequent correct (error) change-of-mind choices. Probability of change-of-mind for subsequent correct choices peak at *ε* = 3.2, before decreasing. (**b**) Response times are slower during change-of-mind (regardless of whether they are correct (dashed green) or error (bold brown)). Error bars are SEM.

**Figure 5.**
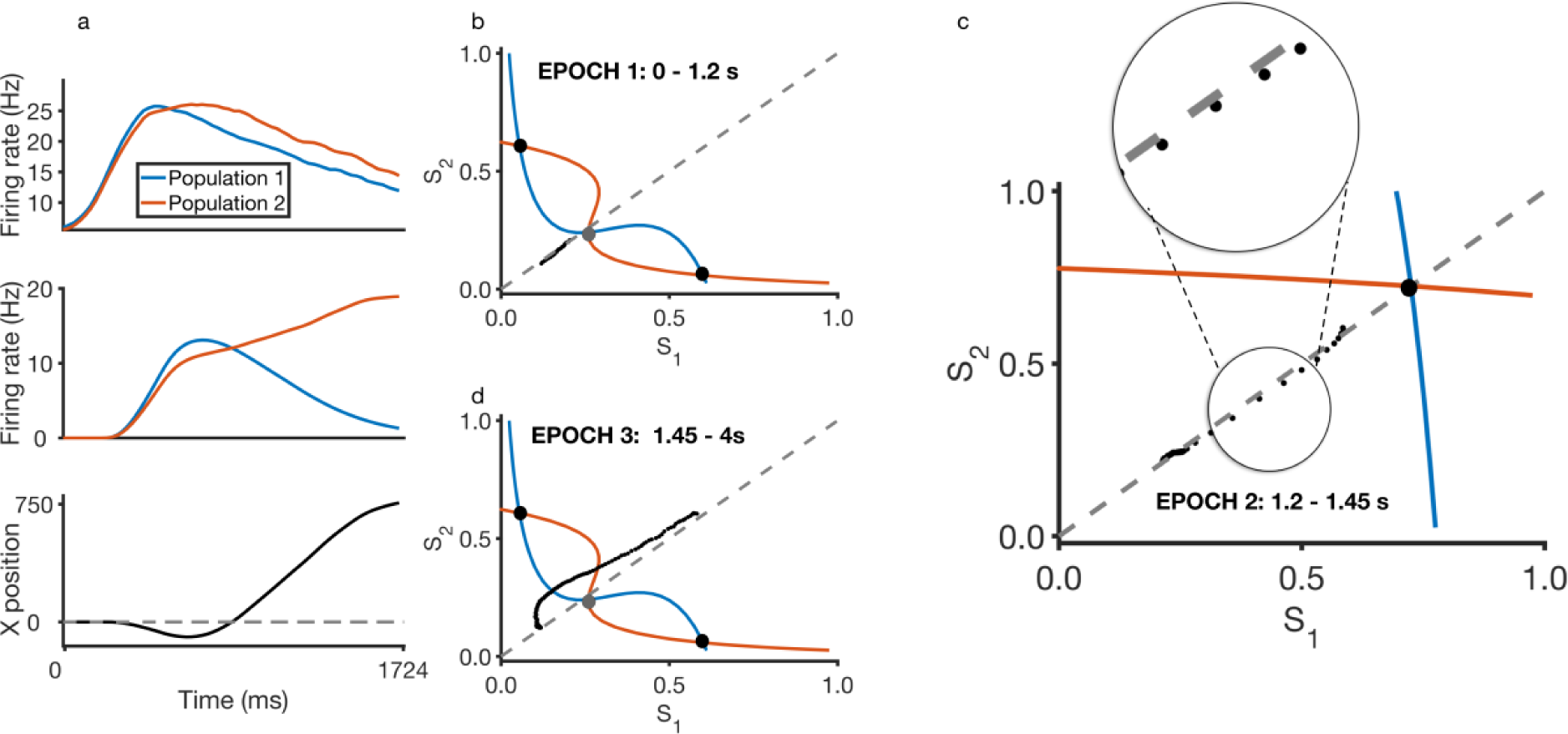
Neural circuit mechanism of change-of-mind behaviour. (**a**) Trial-averaged (n=17) timecourse of firing rates in sensorimotor module (top), motor module (middle) and corresponding motor trajectory (bottom). Evidence quality ε = 3.2 (favouring population/choice 2/Right). Populations compete after stimulus onset (time 0). As motor starts moving in one direction (without reaching the target), a reversal of neural activity dominance in sensorimotor module occurs, leading to a change-of-mind. Note: final decision is made by the motor output in X space (bottom). (**b**) Immediately upon stimulus onset (ε = 3.2, favouring choice 2/Right), the sensorimotor population activity trajectory (black dotted line) in phase space deviates from phase plane diagonal. Black filled circles: stable steady states representing the two choices i.e. choice attractors; grey filled circle: saddle-like unstable steady state. Refer to main manuscript regarding content of the phase plane (e.g. nullclines) (**c**) During the middle epoch of the trial, large excitatory feedback from uncertainty-monitoring module causes phase plane to reconfigure, and a new choice-neutral stable steady state appears which aids the initially losing neural population (population 2). Trajectory is now drawn towards this stable steady state, towards the phase plane diagonal. Inset: Zoom in. (**d**) During the later epoch of the trial, both sensorimotor populations receive lesser excitatory feedback from the uncertainty-monitoring module, resulting in the phase plane reverting closer to the previous condition during the early epoch of the trial.

The general model behaviour, ultimately reported at the motor neuronal populations, is qualitatively similar to the neuronal firing rates and psychophysical (choice accuracy and response time) data observed in two-choice reaction time experiments^4,39,40^. Specifically, the neural activity of the winning (sensorimotor/motor) neuronal population ramps up faster for higher evidence quality (*ε* = 25.6% cf. 3.2%; equivalent to motion coherence in random dot stimulus – see Methods) (Fig. 1b, top and bottom panels); accuracy increases monotonically with evidence quality (Fig. 1c) while reaction time decreases (with error choices slower than correct choices) (Fig. 1d; compared with^39,41^). A choice is considered to be made when one of the activities of the motor neuronal populations crosses a prescribed threshold of 17.4 Hz. The motor neuronal population activity is also directly mapped onto the motor output or positional space (see Methods and below).

Importantly, the (phasic) activity of the uncertainty-encoding neuronal population is higher for trials with higher uncertainty (due to lower evidence quality) (Fig. 1b, middle panel). This rise-and-decay activity around the motor movement onset is consistent with observations from neural recordings in animal and human studies^6,11,25,42^. More specifically, single neuronal firing activity in the orbitofrontal cortex (OFC) (from rodents)^6,11^, EEG^25^ and fMRI^42^ recordings in humans exhibited this rise-and-decay pattern in experimental studies of decision-making under uncertainty (Supplementary Fig. 3), and these activities are higher with higher decision uncertainty. We shall henceforth use this phasic neural activity as an indicator of decision uncertainty monitoring in real-time, and the temporal integral of its neural activity (i.e. area under the curve as a proxy for any downstream neural integrator) as a readout of the decision uncertainty (see Methods). Further, a tonic constant excitatory bias input to the uncertainty-encoding population (Fig. 1a) is required to provide overall excitation (see Methods). As will be shown below, when trials are sequentially dependant (i.e. a reward is only received when a pair of coupled trials result in two correct choices), this same parameter is linearly varied based on the level of uncertainty in the first trial, influencing the uncertainty level (and response time) of the second trial^43^ (see below and Methods).

### Model accounts for relationships among decision uncertainty and psychophysics

We next simulate with our network model to replicate the key experimental findings related to decision uncertainty and confidence as discussed in the Introduction. As most of the decision uncertainty and change-of-mind tasks are based on two-choice reaction-time task paradigms, we shall only focus on such paradigms. Our model first replicates choice accuracy decreasing monotonically with decision uncertainty (Fig. 2a), while producing the ‘<’ pattern^6,11,33,34^ of decision uncertainty (Fig 2b), in which decision uncertainty is higher for lower (higher) evidence quality in correct (error) choices^6,34^ (compared to Supplementary Fig. 1). This pattern also correlates with the response time pattern in Fig. 1d. We further explore this by performing a linear regression (^2^=0.993) on all our simulated response times with decision uncertainty levels (maximum activity, see Methods) and found very strong correlation (Pearson’s r = 0.85) between the two as observed in experiments^12^ (see Supplementary Figure 6).

**Figure 6.**
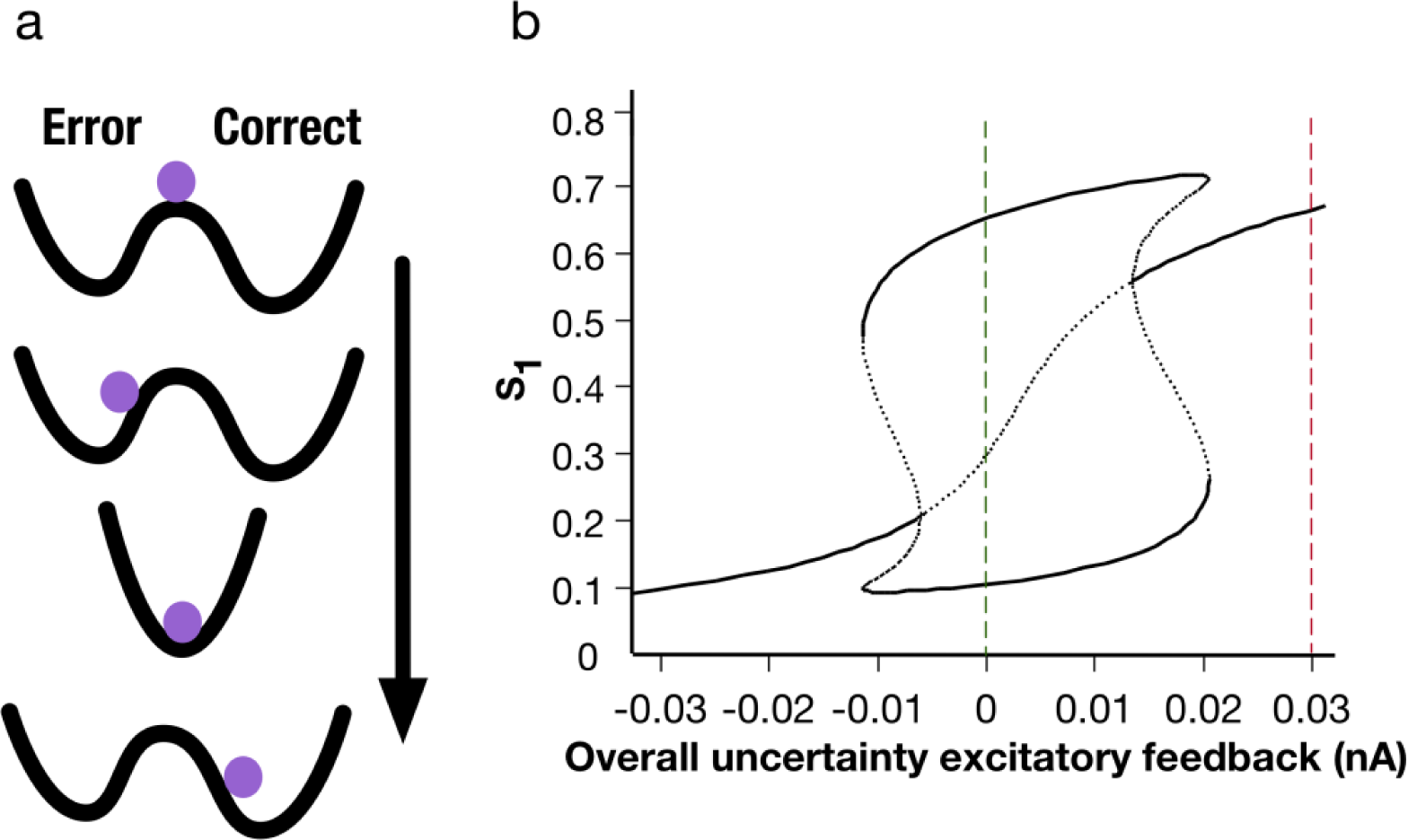
Uncertainty-induced symmetric stable steady state causes change-of-mind. (**a**) Top-to-bottom: Hypothetical “potential well” of network changes over epochs within a trial (arrow). When making a choice between two alternatives, the strength of the stimulus (and noise) drives the ball towards one of the two wells (in this case, an error choice). A transient strong excitatory input (due to excitatory feedback from uncertainty-monitoring module) changes the “energy” landscape into one centralized deep well, allowing a higher chance to change its initial decision. (**b**) Bifurcation (or stability) diagram of the activity of a neuronal population selective to choice 1/Left in the sensorimotor module, S_1_, with respect to variation in the overall excitatory feedback input current from the uncertainty-monitoring module. Evidence quality ε=0. Black bold: stable steady states; black dotted: unstable saddle steady states. Green dashed: initial low uncertainty-induced excitatory feedback and lying within the winner-take-all regime. Red dashed: intermediate epoch of a trial with large uncertainty-induced excitatory feedback – only one stable steady state exists. Later epoch of a trial reverts back to green dashed line.

To explain the results in Figs. 2a and b, we map out the neural activity of the uncertainty-encoding population (denoted by the colours in Figs. 2c and d) with respect to the evidence quality and total input to the uncertainty-encoding neuronal population. Based on Figs. 2c and d, it is clear that as long as the total input is high, and there is sufficient time (i.e. long response time – see Fig. 1d) for the uncertainty-encoding population to integrate its input, the uncertainty level will be high, regardless of correct or error responses. From the perspective of the network dynamics, for correct responses with low evidence quality, the inhibition to the uncertainty-encoding population will initially be higher, i.e. lower total input. This leads to an initial weaker excitatory feedback to the sensorimotor neural populations, causing the ramping-up speed of the latter’s activity to become slower, which in turn results in a prolonged response time. The longer response time allows the uncertainty-encoding population to have more time to integrate and eventually attains a higher activity level, i.e. encodes higher uncertainty. The activities of the competing sensorimotor populations will also eventually deviate (i.e. have a clear winner), resulting in higher total input (i.e. less inhibition) to the uncertainty-encoding population (moving vertically upwards in Fig. 2c, left side). For correct responses with higher evidence quality, the response times are typically faster (Fig. 1d, blue), and hence allowing for less time for the uncertainty-encoding population to integrate, leading to lower uncertainty activity levels (moving vertically upwards in Fig. 2c, right side; see also Supplementary Fig. 7). However, for error responses, the response times are longer for higher evidence quality (Fig. 1d, red), and that allows for more time for the uncertainty-encoding population to integrate. This results in higher uncertainty levels (Fig. 2d, right side). See Supplementary Fig. 8 for a sample trial with long response time where the uncertainty-monitoring module has sufficient time to integrate.

Previous work using a multi-stage decision task paradigm has shown that the level of uncertainty in a decision can affect the response time in a subsequent decision – a form of optimal strategy^43^. Specifically, this only occurs if the reward is tied to two consecutive decisions being answered correctly (i.e. coupled trials). By allowing the same tonic bias input to the uncertainty-encoding population in the second trial to vary linearly based on the decision uncertainty in the first trial of each pair of coupled trials (see Methods), our model can replicate this behaviour (Fig. 3a), exhibiting a prolonged response time in the second decision if the first decision is correct. (Fig. 3a). This trend holds regardless of the evidence quality, with the exception of the easiest difficulty level (due to very low uncertainty levels during these tasks; see Figs. 1d and 2c). The model naturally accounts for this as the neural activity encoding the uncertainty level in the first decision is carried over to the second decision – e.g. higher tonic input with higher decision uncertainty level in the previous trial (Methods). This in turn accelerates or decelerates the ramping-up of neural activity in the sensorimotor populations and hence decreases or increases the response time, respectively.

Next, we sort the simulated trials based on the outcome of both the 1^st^ and 2^nd^ decisions (in each coupled pair) (Fig. 3b), i.e. correct-error combinations. Interestingly, the model predicts a slightly larger difference when the second responses are error choices (red lines) than when the second responses are correct choices (blue lines). This difference (between the correct and error choices) is more pronounced with increasing evidence quality. This can be explained by Fig. 2d: due to the prolonged response time during error choices with higher evidence quality (Fig. 1d) (leading to longer integration time for the uncertainty-encoding population) and a higher total input to the uncertainty-encoding population, higher uncertainty level is reached. Hence, the larger difference.

### Model accounts for change-of-mind behaviours

Previous studies have shown that change-of-mind during decision-making usually leads to the correction of an impending error^4,35^. Although previous studies have linked change-of-mind to the temporal integration of noisy stimulus^4,35^, we demonstrate that the simulated change-of-mind in our biologically-motivated model is due not only to noise, but more importantly, to the necessity of an excitatory feedback mechanism induced by decision uncertainty (Supplementary Figs. 4, 7 and 8). In particular, our network model replicates the observation^4,35^ that the probability of change-of-mind decreases monotonically with evidence quality with the majority of trials leading to ultimately correct choices (Fig. 4a). Further, and consistent with existing observations^4,35^, changes to correct choices peak at an intermediate evidence quality level before gradually decreasing (Fig. 4a) (compared to Supplementary Fig. 3). Moreover, our model predicts that response times are slower during change-of-mind, regardless of evidence quality (Fig. 4b, overlapping bold brown and dashed green lines). When there is no uncertainty excitatory feedback loop, decision uncertainty can still be encoded (Supplementary Fig. 5) but there is no change-of-mind (Supplementary Fig. 4). This suggests that for the biophysically-constrained network model, noisy fluctuation may be necessary but not sufficient to allow significant change-of-mind behaviour. Importantly, a choice-neutral stable steady state (or attractor) due to nonlinearity may be needed.

Experimental observations have shown that the neural instantiation of change-of-mind is associated with a reversal of dominance of neural activities over time within a trial^44^. In our model simulation with change-of-mind, the firing-rate activities of the competing sensorimotor neuronal populations reverse their order of dominance over time within a trial (see Supplementary Figs. 8 and 9a for sample change-of-mind trials). Fig. 5a shows the trial-averaged activity traces of such reversal condition, which can be directly mapped, via the motor neuronal population activity (activity is shown in Fig. 5a, middle panel) into a motor output position in the spatial X direction (Fig. 5a, bottom; see Supplementary Fig. 9a bottom for a sample trial). We can observe switching of neural activity dominance of the sensorimotor neuronal populations before subsequently being dominated by population 2 (selective to rightward choice) (Fig. 5a, top). Note that although the switching of dominance can be small, the difference in activities is integrated and amplified by the motor neuronal populations (Fig 5a, middle), leading to an initial bias towards choice 1/Left (negative X position) (see also Supplementary Figs. 8 and 9a). Further, it should be noted that activities of both sensorimotor neural populations can return to their spontaneous levels – but the activities of the motor neuronal populations could still continue to integrate over time, magnifying the difference in sensory evidence, and hence the motor output can move towards a choice target (Fig. 5a, bottom; see also Supplementary Fig. 9a for a sample trial).

### Neural circuit mechanism of change-of-mind behaviours

Next, we will apply dynamical systems analysis^37^ to demonstrate that this reversal phenomenon is caused not only by noise and strong sensory evidence favouring one population over the other, as indicated in previous modelling work^35^, but also due to the effective excitatory feedback of the uncertainty-monitoring module. Similar to our previous work^37,45^, we plotted the phase planes of the activities of the sensorimotor neuronal populations – which are governed by their slow (NMDA-mediated) population-averaged synaptic gating variables, S_1_ and S_2_ (Figs. 5b-d). These gating variables are monotonic functions of their associated neuronal population firing rates^37,45^. The stimulus is presented with low evidence quality (*ε* = 3.2%). Shown in blue and orange curves in Figs. 5b-d are the nullclines of the sensorimotor module, and their intersections are the steady states – the middle saddle-like steady state (or saddle fixed point) in Figs. 5b and d is unstable while the more off-diagonal ones are stable steady states associated with the choices (or choice attractor states) (Methods). For the latter, the choice attractor closer to the S_1_ (S_2_) axis represents the stable (final) state for making choice 1/Left (2/Right).

With a difficult task (small bias in the phase plane), the sensorimotor neuronal populations integrate sensory evidence and ramp up their activities towards one of the two choice attractors, and on average, almost along the phase-plane diagonal (Fig. 5b, black dotted trajectory). Fluctuations due to noise contribute mainly to the initial dominance in the neural activities, in this case favouring choice 1/Left. This leads to high inhibition of the uncertainty-encoding population and weak excitatory feedback to the sensorimotor populations. The prolonged ramping up of the activities of the sensorimotor populations eventually allows integration of the activity of the uncertainty-encoding neuronal population and provides excitatory feedback to the sensorimotor module. This leads to the reconfiguration of the phase space and the creation of a new central and choice-neutral stable steady state, to which the trajectory of the sensorimotor module activity is now drawn into (Fig. 5c). Notice that the choice attractors have vanished. Furthermore, while the trajectory is being drawn, it moves closer towards and crosses the diagonal line (Fig. 5c). Importantly, the model suggests that this new stable steady state plays an important role in change-of-mind – it provides the initially losing neuronal population a higher chance of winning.

Due to the transient nature of the uncertainty-encoding neuronal population activity (Fig. 1b, middle, and Supplementary Fig. 8), the excitatory feedback returns to baseline level, and the phase plane reverts to its initial configuration (Fig. 5d) (prior to the activation of the uncertainty-monitoring module (Fig. 5b)). This causes the trajectory to move towards the higher part of the phase plane and, coupled with noise, leads to a change-of-mind behaviour. Overall, this is reflected in the reversal of dominance in the neural activities of the motor populations (Fig. 5a, middle) and motor movement (negative-to-positive) direction (Fig. 5a, bottom) (see also Supplementary Fig. 9). It should be noted that in the model, the final decision is determined by whether the firing rate of motor neural populations, which themselves are neural integrators, reach a prescribed target threshold (see Methods). Thus, change-of-mind could still occur even if the activity reversal is not clearly observed in the sensorimotor module.

In our analyses we found that the new central choice-neutral stable steady state is less likely to emerge with higher evidence quality due to shorter response time and weaker excitatory feedback from the uncertainty-monitoring module (Figs. 2c and d; Supplementary Fig. 7). This explains why higher evidence quality generally leads to lower probability of change-of-mind^4,35^ (Figs. 4a, black). For lower evidence quality, the phase plane is almost symmetrical (Fig 5b). Thus, the network is likely to make an error choice initially due to noisy fluctuations. This can lead to longer integration time for the uncertainty-monitoring module and provides stronger excitatory feedback – in the form of a transient, centralized attractor state – and consequently, correcting the decision. Hence, this explains why there are more correct change-of-mind trials than error change-of-mind trials. However, increasing the evidence quality leads to lower probability of change-of-mind, as discussed above. This explains the observed peak in probability of correct changes-of-mind (Fig. 4a and Supplementary Fig. 2).

## Discussion

We have proposed a novel neural circuit computational model that encodes decision uncertainty, the reciprocal of decision confidence. Decision uncertainty in the model can be represented in real-time for online excitatory feedback and for controlling decision dynamics. Our uncertainty-monitoring module was developed based on transient neural dynamics observed in animal and human neuroimaging studies^6,25,42^, and the relationship between choice certainty, evidence and response time^12,33,34^ (e.g. Supplementary Figs. 1-3). Building on our previous decision-making model^37^, our extended neural circuit model can account for several observations commonly found in experimental studies of decision confidence and change-of-mind^4,6,34,35^.

A seminal paper has shown neuronal firing rates from the OFC^6^ can signal decision uncertainty encoded through its phasic activity, as in our model’s uncertainty-encoding population. Specifically, the magnitude of the firing rates in single neuronal recordings in OFC^6^, peaking around the response initiation time. This peak is higher the longer the animal waits before opting out (a measure of decision uncertainty level) (see Supplementary Fig. 3a). This work was extended^11^ by showing that inactivation of OFC neurons during an opt-out waiting task causally affected the animal opting out (i.e. decision uncertainty reporting) behaviour. More recently, EEG (theta band) and fMRI recordings have also shown neural activity exhibiting similar characteristics^25,42^, with phasic activity peaking around the response initiation, and the peak was higher with higher reported uncertainty or when an error was detected by the participants.

We have proposed a model that was able to exhibit higher levels of decision uncertainty and lower choice accuracy with more difficult tasks ^6,10,36^ (Fig. 2a). Further, the model showed higher decision uncertainty with lower evidence quality for correct choices, but counter-intuitively, lower decision uncertainty with higher evidence quality for incorrect choices, in line with the previously observed ‘<’ pattern^6,11,33,34^ (Fig. 2b, Supplementary Fig. 1). This was explained by the faster response times for correct choices, with lesser integration time for the uncertainty-monitoring module, which led to lower decision uncertainty (Figs. 2c and d). For error choices, the integration time was longer with higher evidence quality (Fig. 1d). This led to longer integration time for the uncertainty-monitoring module and hence higher decision uncertainty. Furthermore, the uncertainty-monitoring module provided a closed-loop recurrent network mechanism of excitatory feedback with the sensorimotor neuronal population, enhancing the latter’s responses. This was reminiscent of a dynamic gain or urgency mechanism^46,47^. Future work could test this mechanism, e.g. using a task paradigm that produces fast error choices^48^ and determining whether the ‘<’ pattern is absent.

By utilising a proxy memory mechanism instantiated in the existing tonic bias input to the uncertainty-encoding neural population, our model was also able to show that decision uncertainty from a correct first trial caused a slower response time in the second trial, compared to when the first trial was incorrect (Fig. 3a). Moreover, the model predicted a slightly larger difference in response times when the second responses were error choices than when the second responses were correct choices (Fig. 3b). This difference was more pronounced with increasing evidence quality. Future work could test our model’s prediction, for instance, by direct micro-stimulation or inactivation of the uncertainty-encoding (or outcome anticipation) neurons in the medial frontal cortex e.g. OFC in rodents^6,11^ or subregions in the human frontal cortex^49^.

It should be noted that the above multi-stage decision paradigm is a special case of sequential decision-making^43^. Specifically, two coupled decisions have to be correct in order to receive a reward. The time delay from the first choice to the next stimulus onset (response-stimulus interval, RSI) is sampled from a truncated exponential distribution (range 0.3–1.0 s; mean 0.57 s). When simulating this paradigm, we have reset the uncertainty bias upon the completion of every pair of coupled trials. Thus, our implementation of the paradigm would not be affected by the RSI. Our stored uncertainty bias could perhaps be allowed to decay over time, for instance, similar to our previous work^50^. However, to the best of our knowledge, this multi-stage decision study^43^ is the only published work that links decision confidence to response times with sequential dependency, and we defer such speculation to further experimental evidence.

The results in Figs. 3a and b could be explained by the uncertainty level mappings (Figs. 2b, c and d). Specifically, in pairs of coupled trials, errors in first decisions led to a higher tonic bias input (and subsequently, higher overall input, Fig. 2d) in second decisions, due to higher uncertainty levels in first error decisions (Fig. 2b, red) than correct decisions (Fig. 2b, blue), which resulted in stronger excitatory feedback to the sensorimotor module. This led to faster activity ramping up of the sensorimotor populations, which in turn caused faster error (than correct) response times in second decisions. Furthermore, Fig. 3b showed that such differential effect would be more prominent for higher evidence quality.

The same model could exhibit changes-of-mind which were more likely to occur with lower evidence quality^4,35^ (Fig. 4a, black). Specifically, the model showed that changes-of-mind were more often accompanied by correcting an impending error choice – hence more error-to-correct changes than correct-to-error changes (Fig. 4a, green vs red), consistent with previous observations^4,35^. Furthermore, the likelihood of error-to-correct changes slightly peaked at an intermediate level of evidence quality before decreasing as the task becomes easier^4,35^ (Fig. 4a, green). The model predicted slower response times during changes-of-mind, regardless of evidence quality (Fig. 4b). Critically, when we removed the excitatory feedback from the uncertainty-monitoring module to the sensorimotor module, the decision uncertainty could still be encoded, but there was no change-of-mind (Supplementary Figs. 4 and 5). This demonstrated the importance of the uncertainty-induced excitatory feedback on changes-of-mind.

We used phase-plane analysis to explain the change-of-mind phenomenon. First, the process of change-of-mind could be understood in terms of the sensorimotor network state being attracted to three distinct basins of attraction: the initial choice, then to the central choice-neutral ‘uncertain’ state, and finally to the other choice. With higher evidence quality, we found that the correct choice attractor dominated the phase plane, with its generally larger basin of attraction (e.g. Supplementary Fig. 10; see also^37^) and the central attractor was less likely to appear due to the weaker uncertainty-based excitatory feedback (e.g. compare Supplementary Fig. 7 to Supplementary Fig. 8). This explains the monotonic decrease of the probability of change-of-mind (Fig. 4a). In other words, changes-of-mind did not occur due to the heavily biased phase plane and fast response times. However, at low evidence quality levels (ε < 4%), the phase plane was almost symmetric (Fig. 5b), which led to more initial errors (Fig. 4a). Under such low evidence quality, it was increasingly likely that the network would make an initial error choice^37^. This led to longer integration time of the decision uncertainty-monitoring module and provided stronger excitatory feedback – in the form of a transient, central choice-neutral stable steady state – and eventually, correcting the decision (Figs. 5c and d, and Supplementary Figs. 8 and 9). On the contrary, increasing the evidence quality led to lower probability of changes-of-mind. This explains the peak in probability of correct changes-of-mind at an intermediate evidence quality (Fig. 4a; Supplementary Fig. 2). The model further suggested that during changes-of-mind, noisy fluctuation around the phase-plane diagonal led to subtle deviations early in the trial (Fig. 5). The downstream motor module, which was itself a neural integrator, amplified any slight deviation and led to movement being initiated towards a choice target (Figs. 5a, and Supplementary Figs. 8 and 9).

Fig. 6a illustrates a hypothetical decision ‘potential well’^37^ that summarizes our key findings for change-of-mind– the central attractor, caused by the excitatory feedback from the decision uncertainty-monitoring module, and coupled with noise, can allow an initial choice to be altered. The strength and duration of this attractor depends on the evidence (and elapsed time) for temporal integration by the sensorimotor neuronal populations.

To provide further insights, we have provided a bifurcation (or stability) analysis of the activity of a neuronal population (selective to choice 1/Left) in the sensorimotor module, S_1_, with respect to the systematic variation (bifurcation parameter) of the overall excitatory feedback input current from the uncertainty module with evidence quality *ε* = 0 (Fig. 6b). The stable steady states are denoted by black lines, while dotted lines represent the unstable saddle steady states. During the initial epoch of a trial, this excitatory feedback input from the uncertainty-monitoring module (specifically the uncertainty-encoding neuronal population) to the sensorimotor population is very low or zero (green vertical dashed line). This is the regular winner-take-all regime^37^. As sensory evidence is accumulated in the sensorimotor populations, the uncertainty level is increased, which leads to higher excitatory feedback from the uncertainty-monitoring module. When the overall excitatory feedback is sufficiently large (larger than ~0.03nA in our simulations (vertical red dashed line)), the network is attracted towards the only choice-neutral stable steady state. However, this effect is only transient – in a later epoch of a trial, the neural activity of the uncertainty-encoding neuronal population may return towards a lower level, and the decision network would once again return to the winner-take-all regime^37^ (vertical green dashed line).

Unlike previous neurocomputational models^35,51^, our model does not rely on explicitly reversing the stimulus input to neural populations or having a relatively low (first) decision threshold (to induce faster errors). Further, it does not rely on abstract mathematical calculation of decision uncertainty^28^. Inspired by neural evidence of decision confidence^6,25^, we have a dedicated neural module that has a plausible circuit architecture resembling a cortical column that monitors and quantifies the decision uncertainty and controls decision dynamics via feedback.

Our model complements simpler computational cognitive models such as the extended drift-diffusion models^4,52,53^, by providing a neural circuit perspective on the neural mechanism behind decision confidence/uncertainty and change-of-mind. Specifically, our model links to psychophysical data (Figs. 1c, 1d, 2a, 2b, 3a, and 4a) and also directly relates to neurophysiological data (Figs. 1b and 5a, and Supplementary Figs. 7, 8 and 9a), which simpler models cannot readily do. Hence, both psychophysical (Figs. 3b and 4b) and neural (Figs. 1 and 5, and Supplementary Figs. 7-10) predictions are naturally embedded in the model. That said, such biologically-motivated (mean-field) models can be linked back (through various model reductions and assumptions) to simpler cognitive models such as the drift-diffusion models^37,45,53^.

Several cognitive models have been proposed to model different roles of the medial prefrontal cortex (mPFC) and anterior cingulate cortex (ACC), which include error prediction^54^. However, it is unclear how the predicted ACC signals (i.e. negative and positive ‘surprise’ signals) in such models can influence the dynamics of decision formation^55^. in addition to explicitly modelling the dynamics of perceptual decision uncertainty, our model provides an account of the effect of decision uncertainty on the dynamics of the decision formation process; from sensory evidence integration up to motor output. This results in decision changes ‘on the fly’ leading to change-of-mind within a trial.

Our distributed neural circuit model is more realistic than other biologically-motivated computational models of decision confidence or change-of-mind^35,51^. Evidence shows perceptual decisions are performed and distributed across multiple brain regions^56^. Specifically, the activity of our motor module can be directly transformed to motor positional coordinates, hence directly maps to physical output. Our model with the feedforward neural integrator architecture (from sensorimotor to motor module) suggests that the reversal of neural activities resembling a change-of-mind could be more clearly identified in more motor-based neurons than sensory-based neurons (Fig. 5 and Supplementary Figs. 8 and 9). Future experiments could show the difference in neural dynamics in different brain regions during change-of-mind tasks, e.g. via dual recordings at the sensory and motor-based brain regions. Importantly, we consider our proposed model to be a reconciliation of two views in the literature on how changes-of-mind can occur: In our model, bottom-up evidence^4,57^ is continually accumulated after the choice is made through recurrent excitation and noise fluctuation, while top-down^25,42^ evidence is accumulated through the excitatory feedback loop (from the uncertainty-encoding population).

In summary, our work has provided a neural circuit model that can compute decision confidence or uncertainty within and across trials while also occasionally exhibiting changes-of-mind. The model can replicate several important observations of decision confidence and change-of-mind and is sufficiently simple to allow rigorous understanding of its mechanisms. Taken together, our modelling work has shed light on the neural circuit mechanisms underlying decision confidence and change-of-mind.

## Methods

### Psychometric and chronometric function

We used a Weibull function^58^ to fit the psychometric function *p* = 1 − 0.5 *exp*(−*ε*/*α*)^*β*^, where *p* is the probability of a correct choice, *ε* is the evidence quality, which, in the case of the random-dot stimulus^59,60^, is equal to the motion coherence level (*c′*). With the parameters used with our model (see Supplementary Table 1), *α* (the threshold at which the performance is 85%) is set to 7.32%, while *β*, the slope, is equal to 1.32. We defined the model’s response (or reaction) time as the overall time it took for the motor neuronal population activity to reach a threshold value of 17.4 Hz (mimicking a motor output and physically reaching a choice target) from stimulus onset time. This is equivalent to the motor position reaching some threshold value (see below).

### Modelling sensorimotor populations using two-variable model

We used the reduced version of the spiking neural network model^51^ described by its two slowest dynamical variables, which are the population-averaged NMDA-mediated synaptic gating variables^37^. The dynamics of the two neuronal populations can be described by:

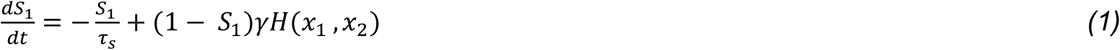

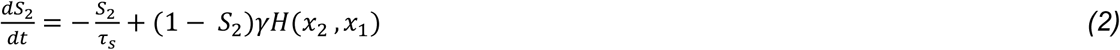

where the two excitatory neuronal populations representing the two choice options are labelled 1 and 2, and the *S*’s are the population-averaged NMDA-mediated synaptic gating variables. *γ* is some fitting constant based on previous work^37^. *τ*_*s*_ denotes the synaptic gating time constant (100 ms) constrained by NMDA receptor physiology. *H* denotes the nonlinear single-cell input-output function fitted to that of a spiking neuronal model. The firing rates of the sensorimotor populations can be described by these three equations:

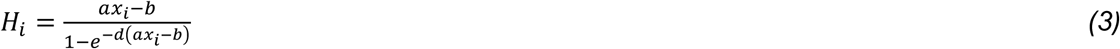

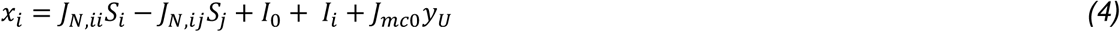

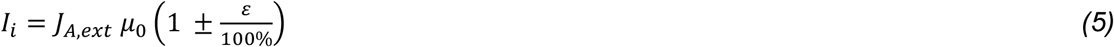

where *a*, *b*, *d* are parameters for the input-output function fitted to a leaky integrate-and-fire neuronal model^37^. The dynamical variables S_i_ and S_j_ are from Eqs. 1 and 2. *J*_*N,ii*_ and *J*_*N,ij*_ are synaptic coupling constants from recurrent connections. *I*_0_ denotes a constant value that represents an effective bias input from the rest of the brain. *I*_i_ denotes the excitatory stimulus input to population *i*, and is proportional to the evidence quality *ε*, with the stimulus strength constant denoted by 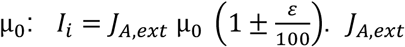. *J*_*A,ext*_ represents the external synaptic coupling constant. In addition to the features in the previous work^37^, the strength of excitatory feedback from the uncertainty-encoding population is controlled by *J*_*mc*0_. Hence, decision uncertainty is monitored and fed back to the sensorimotor populations via excitatory feedback.

### Uncertainty monitoring neuronal populations

A key aim of our modelling work is to understand how the neural circuit dynamics and choice behaviour can be modulated by decision uncertainty. In particular, our proposed uncertainty-monitoring module can lead to error correction through change-of-mind within a trial. It should be noted that, while modelling decision uncertainty, the model was constrained by the neural profile of uncertainty-encoding neurons (or brain regions) observed in experiments (single neuronal recording^6^, EEG recordings^25^, and fMRI recordings^42^).

Two neural populations mimicking a canonical cortical microcircuit were implemented. One population, an inhibitory population, integrates the summed output of the two sensorimotor neuronal populations while another population, and excitatory population, monitors decision uncertainty. Their dynamics are described by:

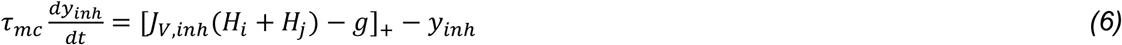

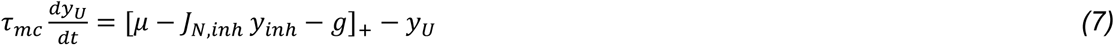

where y_inh_ and y_U_ are the dynamical variables of the inhibitory neuronal population and uncertainty-encoding population, respectively. []_+_ denotes a threshold-linear input-output function (with threshold of 0), with its input argument in units of nA. *J*_*V,inh*_ denotes a synaptic coupling constant from the sensorimotor populations to the inhibitory neuronal population. *H*_*i*_ and *H*_*j*_ are the neuronal population firing rates from the sensorimotor populations *i,j.g* represents some top-down inhibition (1000 nA) on the uncertainty-encoding (and inhibitory) population from beginning of trial, which is removed 500ms from Eqs. (6) and (7) after stimulus onset, respectively (see Supplementary Figure 11 where the effect of this timing feature on the model performance was explored). We used these delay values for all the figures in the main text and in Supplementary Information, unless noted otherwise (see Fig. 1). When the activity of one of the sensorimotor neuronal populations crosses a threshold value (35.5 *Hz*), *g* is reactivated (3000 nA). This results in the activity pattern of uncertainty-monitoring module to mimic data observed in neural recordings^6,25^ (see Fig. 1b, middle panel). *J*_*N,inh*_ denotes the inhibition strength from the inhibitory neuronal population to uncertainty neuronal populations, while *μ* is some excitatory constant bias input that can be modulated (only in multi-stage decisions) by decision uncertainty from the first trial in a pair of coupled-trials (see below).

### Justification of our modelling choices

a) **Tonic activity:** In the uncertainty-encoding population, the tonic activity (i.e. elevated baseline background activity) provides a mechanism by which inhibitory input/signal can be transmitted. In particular, if the receiving neuronal population firing rate was to be silent (i.e. no tonic activity), then only excitatory input can be transmitted. Specifically, in such cases, inhibitory input cannot be transmitted due to the ‘flooring’ effect (i.e. the neuronal firing rate cannot be negative).

b) **Inhibitory-excitatory pair of populations in uncertainty-monitoring module:** The pair of excitatory and inhibitory neural populations in the uncertainty-monitoring module resembles the simplest plausible computational model of a cortical column^38^ – we assume that uncertainty is encoded in the cortex, e.g. the frontal cortex^6,25,42^. Second, and importantly, the key computational role of this inhibitory population is not only to restrain but to modulate the activity levels of the uncertainty-encoding population, and hence influencing the encoding of decision uncertainty (see Results section).

c) **Top-down (dis)inhibition**: Used in our model, such an inhibitory mechanism has been proposed to originate from various brain regions, e.g. involving the superior colliculus and basal ganglia. For example, the threshold crossing (response threshold in our model, which triggers top-down inhibition) could be detected by the superior colliculus via basal ganglia^61,62^. More complex gating pathways in the brain, including disinhibitory circuits, have been proposed that also involve subcortical structures, such as the basal ganglia and thalamus^63^. It should be noted that providing an explicit account of such complex neural circuit dynamics is beyond the scope of this work. As a proxy to modelling such complex and extended neural networks, we have instead modelled just the onset and offset of the top-down inhibition. The onset of this top-down inhibition is assumed to have been learned e.g. through the basal ganglia^64^, which is mediated through neuromodulators.

### Motor neuronal populations

Similar to the uncertainty-monitoring neuronal populations, we dynamically modelled the motor output module using threshold-linear functions (with threshold value of 0). Two neural populations selective for right and left – with mutual inhibition – were used. The persistent activity can be maintained using mutual inhibition to create a line attractor model^65^. The dynamics of the neuronal populations for the two choices (1 and 2) are described by:

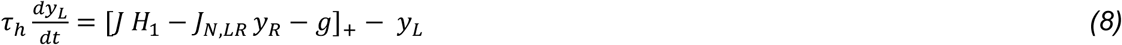

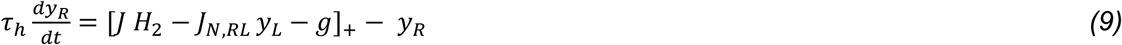

where y_R_ and y_L_ are the dynamical variables of the left and right motor neuronal populations, populations (Fig. 1), and the associated coupling constant J = 1 *nA Hz^−1^*. *J*_*N,ij*_ denotes a coupling constant from population *i* to population *J*. The negative sign indicates connectivity is effectively inhibitory. As for the uncertainty-monitoring module, *g* represents some top-down inhibition (1000 nA) on the motor populations from beginning of trial and is removed when the activity of one of the sensorimotor neuronal populations crosses a threshold value (35.5 *Hz*).

### Mapping the activity of the motor module to X position

The motor module output as a position in the *x* directional space is approximated by a linear function:

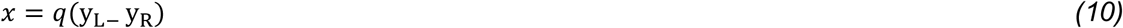

where *q* is a constant scaling factor with a value determined by the equation:

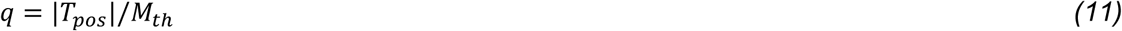

where *T*_*pos*_ is the hypothetical position of the two opposing choice targets. 1366×768 is one of the most commonly used screen resolutions. Therefore, in the model, this value is set to 750 or −750 (close to the edge of the x dimension). *M*_*th*_ is the motor target threshold, set to 17.4 Hz.

### Uncertainty within a single trial

We used two measures to quantify the level of decision uncertainty in a trial. For the first measure, we used the maximum firing rate value of the uncertainty-encoding population for each trial *n*, allowing real-time monitoring of decision uncertainty. For a specific evidence quality value, we calculated the trial-averaged and SEM of these maximal values. For the second measure, we calculated the area under the curve of the firing rate activity over time of the uncertainty-encoding population using the trapezoidal numerical integration scheme for each trial *n*. This provides an overall quantification of decision confidence after a choice is made. It also acts as a proxy for any downstream neural integrator that temporally integrates real-time decision uncertainty information. Again, for each evidence quality value, we calculated the mean and SEM of the areas. Either measure of decision uncertainty is then normalised using feature scaling to bring all values within the range [0,1]. This is done by:

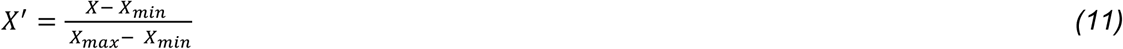

### Uncertainty across coupled trials

In coupled trials, the evidence quality of the second trial was selected probabilistically from a uniform distribution, where *ε* ∈ [0,3.2,6.4,12.8,25.6,51.2]. The area under the curve of the summed activity of the uncertainty-encoding population at trial *n*, *X*_*n*_, is transformed and stored into some activity measure *C* in the subsequent trial. We used a simple linear transformation described by

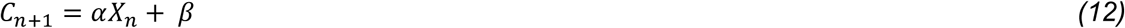

where *n* denotes the trial number, *α* and *β* are scaling parameters. The parameter values set in this work are *α* = 0.008 *nA* and *β* = 0.5 *nA*. This value of *C*_*n+1*_ is then used to modulate the tonic input (and hence baseline activity) of the uncertainty-encoding population (*μ*, in equation (7)) in the second trial using the following update:

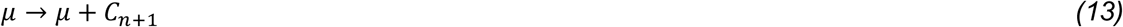

Upon the completion of a pair of coupled trials, the uncertainty bias *C*_*n+1*_, stored in *μ*, is reset to 0.

### Regression and classification of model outputs

We used a smoothing spline function in MATLAB to fit the model’s decision accuracy as function of uncertainty level. We also performed a linear regression on all our simulated response times with decision uncertainty levels (*R*^2^=0.993). The two variables were highly correlated (Pearson’s r = 0.85, p-value = 0) (see Supplementary Fig. 6). The model behaviour is identified to have a change-of-mind if there is a reversal in the order of dominance between the two motor neuronal population firing rates, i.e. if there is a change in the sign of *x* (eq. 10), and a choice target is eventually reached (before a 4 s timeout – see below) (see Fig. 5a, and Supplementary Figs. 8 and 9).

### Simulation and analysis

The code to simulate the model was written in MATLAB (version 2018a) and was run on a Mac OS X workstation. The forward Euler-Maruyama numerical integration scheme with an integration time step of 0.1 ms was used for numerical integration of the dynamical equations (describing dynamics within a trial). Smaller time steps were checked (e.g. 0.01 ms) without affecting the results. XPPAUT^66^ was used to perform dynamical systems (phase-plane) analysis and for parameter search on each neural module and for the bifurcation analysis. The model’s parameter values are summarized in Supplementary Table 1. The model was simulated under a response-time task paradigm with a timeout of 4 s. The stimulus appeared 900 ms after a trial has begun. Only 2.2% of the total simulated trials (8000 trials per condition) were indecision trials in which the motor activity did not cross the 17.4 Hz threshold, i.e. choice target was not reached. These simulated trials were discarded and not included in our analyses.

### Selection of parameter values

Please refer to Table 1 in Supplementary Information for more information on how the parameter values were selected. In some cases, parameters were adopted from previous work^37^. Some parameter values,such as *j*_*mc*0_ (coupling strength between the uncertainty-encoding and sensorimotor populations) and the integration timing parameters were selected to fit qualitative aspects of existing observations (< pattern^6,11,33,34^, probability of changes-of-mind^4^, neural profile of experimentally-observed uncertainty-encoding neurons and regions^6,25^). We describe the effect of changing these parameter values on model behaviour in Supplementary Figs. 4 and 11.

### Code availability

MATLAB and XPPAUT codes were used to simulate the model and generate the figures. They can be found at the following GitHub repository (repo): https://github.com/nidstigator/uncertainty_com_modelling The accompanied README file includes detailed instructions on how to reproduce these figures.

## Acknowledgements

We thank D. O’Hora and A. Zgonnikov for helpful discussions. We also thank P.T. Piiroinen, J. Zhang, S.P. Kelly and R.G. O’Connell for insightful discussions during the early phase of the project, and W.P. Boyce and A. Joshi for critical reading of the manuscript. We appreciate constructive comments from the anonymous reviewers. This work was supported by the Ulster University Vice-Chancellor’s Research Scholarship Award (N.A.), Northern Ireland Functional Brain Mapping Project 1303/101154803 funded by InvestNI and the Ulster University (G.P. and K.W.-L.), and The Royal Society International Exchange Scheme under grant IE151293 (I.R. and K.W.-L.). K.W.-L. was further supported by BBSRC (BB/P003427/1), COST Action Open Multiscale Systems Medicine (OpenMultiMed) supported by COST (European Cooperation in Science and Technology), and The Royal Society – NSFC International Exchanges under grant IE141307.

Author contributions
N.A. and K.W-L. formulated the idea. N.A. and K.W.-L. designed the experiments. N.A. performed and analyzed most of the experiments. N.A. and K.W.-L. analyzed the data. I.R. and G.P. provided additional suggestions relevant to the experiments and analysis. N.A. and K.W.-L. wrote the original draft of the paper. All authors reviewed and edited the paper.

