## Supplementary Information for "Neural Circuit Mechanism of Decision Uncertainty and Change-of-Mind"

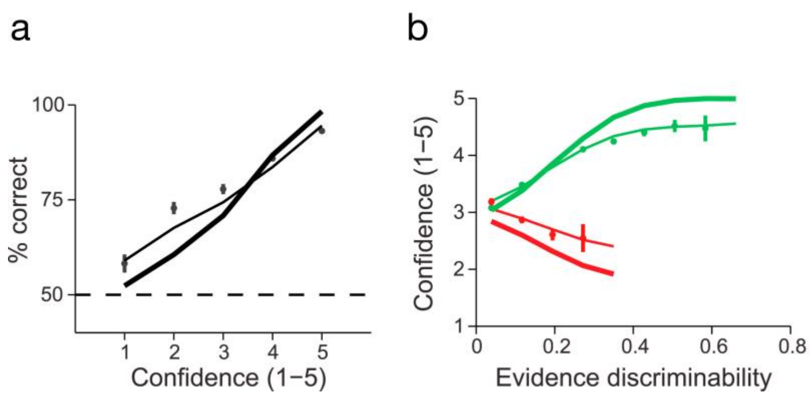


**Supplementary Figure 1. Common characteristics of decision confidence.** (**a-b**) Thick and thin lines are two statistical model fits of an experimental study. Combined data of all subjects (n = 5) (**a**) Choice accuracy as a function of decision confidence. Monotonic increase in accuracy (% correct) with increasing confidence level. (**b**) < (or sometimes called X) pattern of decision confidence with respect to increasing evidence discriminability (quality) i.e. decreasing task difficulty. Decreasing (increasing) confidence for error (correct) choices with increasing stimulus strength. Green: Correct choices; red: error choices. Adopted from[1]. Similar pattern reported in[2–4].

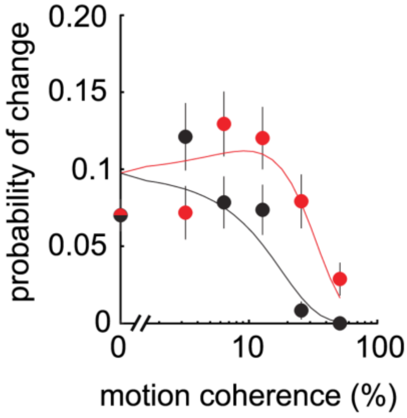


**Supplementary Figure 2. Common characteristics of change-of-mind.** Correct changes-of-mind (red) are generally more frequent than error changes-of-mind (black) and peak at an intermediate task difficulty level (the task difficulty is the % of motion coherence in random dot kinematogram of a motion discrimination task) before decreasing. Data fitted with an extended drift-diffusion model. Adopted from[5].

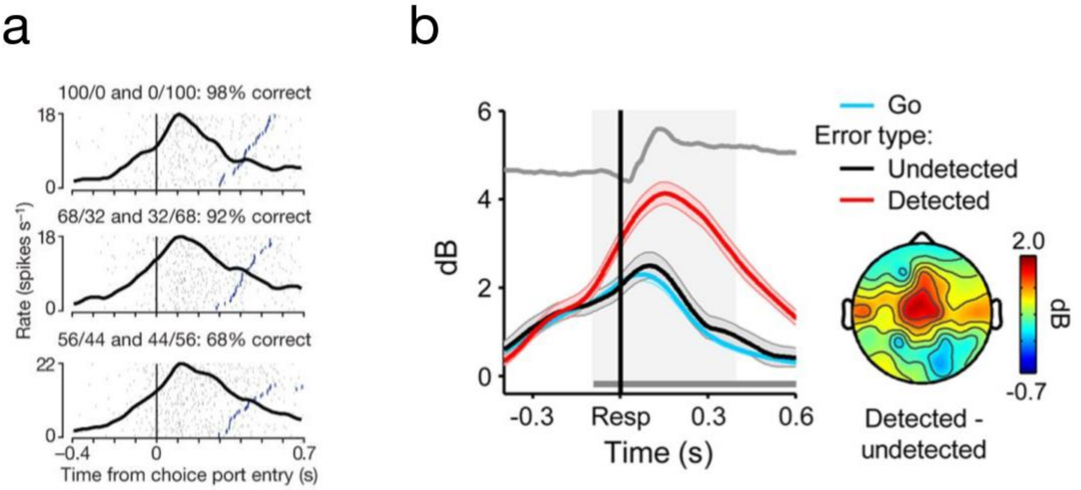


**Supplementary Figure 3. Phasic neural activities of decision uncertainty.** (**a**) Opt-out waiting task in which the animal can opt-out for less-but-sure reward. The time it takes the animal to opt-out, a measure of decision uncertainty, correlates with the single neuronal firing rate in the orbitofrontal cortex (OFC). Each row corresponds to a stimulus difficulty (top: easiest; middle: easy; bottom: difficult). Note the small but non-zero firing activity at baseline, and the slightly higher peak of the firing rate with lower accuracy (bottom row). Adopted from[2]. (**b**) A Go/No-Go decision-making task in which participants can make mistakes. Averaged EEG responses (theta band) (left) from the frontoparietal cortex (right) is associated with participants detecting their errors. Scalp topography shows the distribution of error detection effect (signal-to-noise ratio is shown in dB). Adopted from[6].

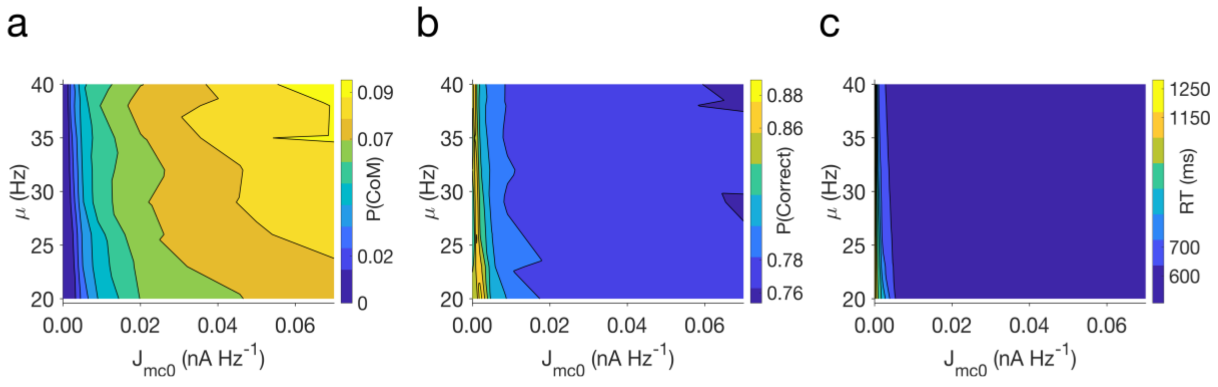


**Supplementary Figure 4. Model performance determined mainly by the strength of the excitatory feedback from uncertainty.** All changes in parameters are simulated with 8000 trials under one condition ($\varepsilon = 6.4\%$). $\mu$: tonic input to the uncertainty-encoding neural population; $J_{mc0}$: excitatory feedback strength from uncertainty-monitoring module to sensorimotor module. Effects across other evidence quality levels can be inferred from the main simulation results (see Fig. 1 and Fig. 4 in the main manuscript). (**a**) Probability of changes-of-mind (CoM), P(CoM). P(CoM) is increased with increasing excitatory feedback strength. (**b**) Accuracy (probability of correct, P(Correct)) increases as the excitatory feedback strength from the uncertainty-monitoring module to sensorimotor module is increased. Saturation at around 77% accuracy due to reaching the fixed decision threshold (35.5Hz, see Methods). (**c**) Faster response times (RTs) are observed when excitatory feedback strength is increased. Saturation of RTs at 600ms due to reaching the fixed decision threshold (35.5 Hz, see Methods).

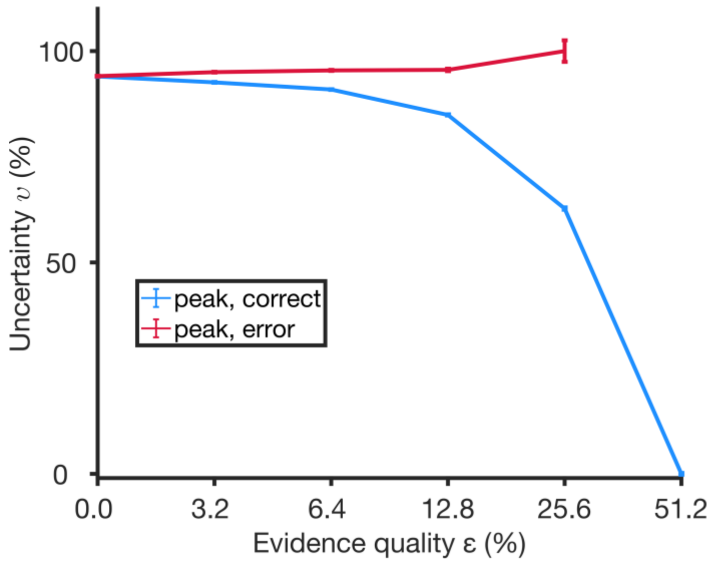


**Supplementary Figure 5. Decision uncertainty as a function of evidence quality from model simulation without the excitatory feedback loop from the uncertainty module**. 8000 trials per condition of evidence quality. List of parameter values used in this simulation can be found in Supplementary Table 1, with the exception of the excitatory feedback strength ($J_{mc0}$) set to 0. Model can encode decision uncertainty without the feedback loop but with no change-of-mind (see Supplementary Figure 4).

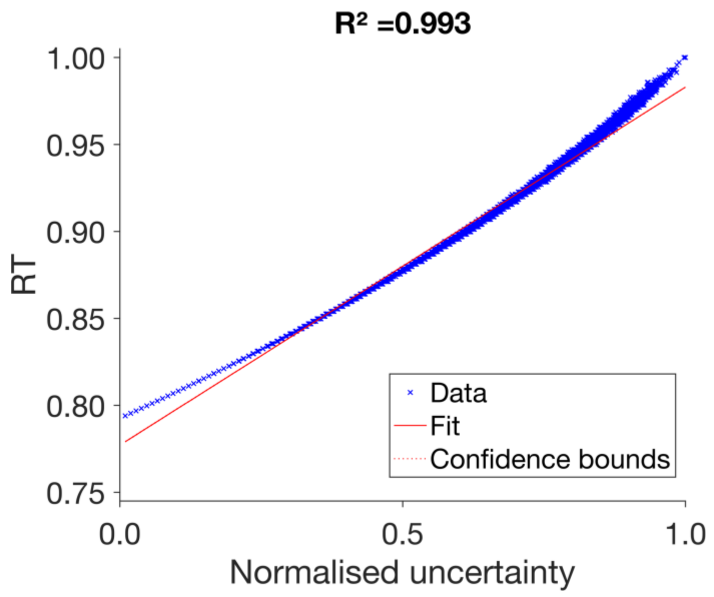


**Supplementary Figure 6. Strong correlation between decision uncertainty level and response time.** Data points (blue) of individual response times (RTs) were obtained from 8000 simulated trials per evidence quality level (48,000 trials). Data indiscriminately consisted of correct, error, CoM and non-CoM trials. Fit (red line) was performed using a linear regression of the uncertainty level as a function of RTs. The two variables (uncertainty level and RT) have a very high (Pearson's) correlation coefficient of 0.85 (p-value = 0) (not shown). Decision uncertainty was calculated using the maximum activity level (similar results with area under the curve method – not shown) (see Methods for further detail). High evidence quality trials that resulted in insignificant change (i.e. <1 Hz) in decision uncertainty levels were excluded from the analysis (6042 trials).

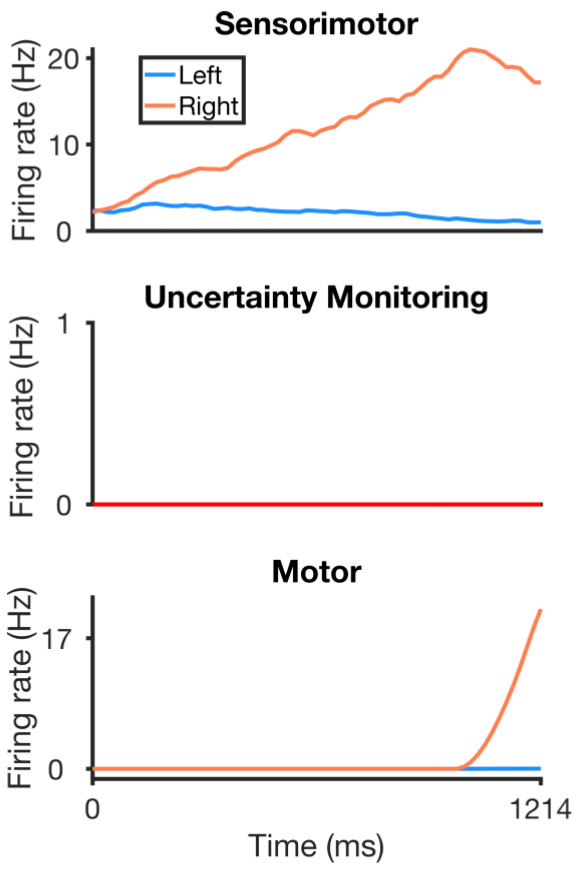


**Supplementary Figure 7. Easy trial: a sample activity timecourse.** Sample timecourse of firing rates in sensorimotor module (top panel), uncertainty-encoding population (middle panel), motor module (bottom panel). Easiest difficulty ($\varepsilon = 51.2$). When the motor activity crosses the 17 Hz threshold, the motor output is assumed to reach a choice target. Due to faster ramping up of activity, the response threshold in the sensorimotor population (35.5 Hz) is crossed before the temporal integration of uncertainty-encoding population begins. Trial completion time (time target threshold was reached) at 1214 ms. Neural population firing rates were calculated by averaging over a time window of 50 ms, slided with a time step of 5 ms.

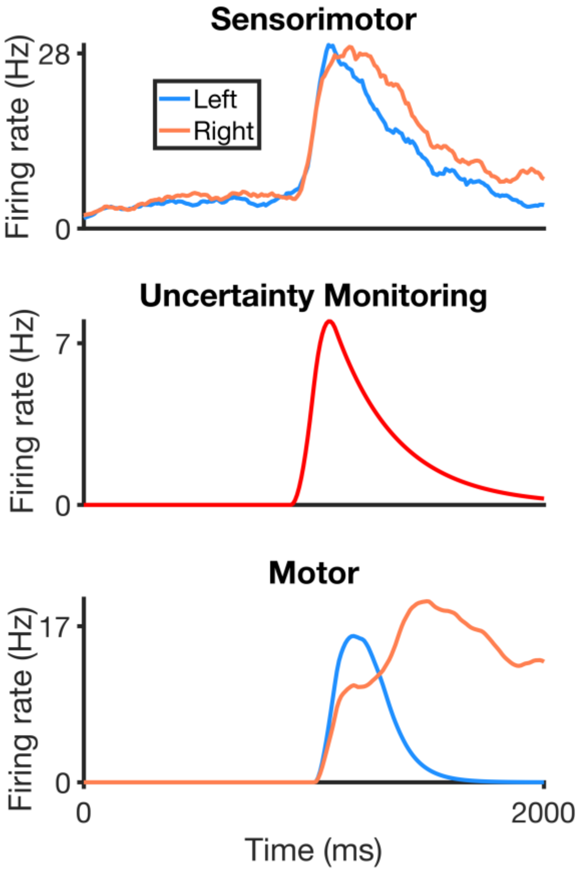


**Supplementary Figure 8. A sample neural activity timecourse during change-of-mind.** Sample timecourse of firing rates in sensorimotor module (top panel), uncertainty-encoding population (middle panel), motor module (bottom panel). ($\varepsilon = 3.2$) exhibiting change-of-mind. Despite the small difference in the dominance of activity in the sensorimotor module, the motor populations continue to integrate over time, amplifying this difference. Trial completion time (time target threshold was reached) at 1637 ms. Neural population firing rates were calculated by averaging over a time window of 50 ms, slided with a time step of 5 ms. Integration is shown until 2000ms to reveal the full dynamics (the phasic nature) of the uncertainty-encoding population activity (middle panel).

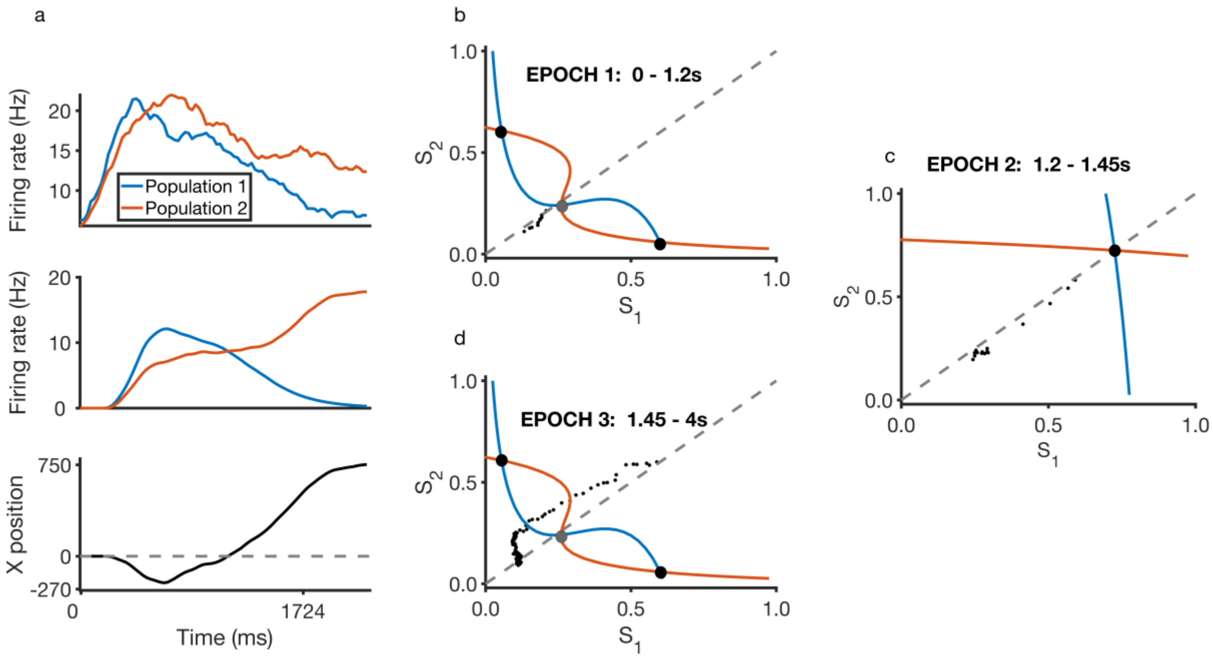


**Supplementary Figure 9. Dynamics of a sample change-of-mind trial.** (**a**) Timecourse of firing rates in sensorimotor module (top panel), motor module (middle panel) and motor trajectory (bottom panel) with bias input (evidence quality) $\varepsilon = 3.2$ (favouring population/choice 2/Right). Left (blue line) and right (orange line) populations compete after stimulus onset. As motor starts moving in one direction (without reaching the target), a reversal of neural activity dominance in sensorimotor module and motor module occurs. This leads to a change-of-mind. (**b**) Immediately upon stimulus onset with evidence quality $\varepsilon = 3.2$ (favouring choice 2/Right), the sensorimotor population activity trajectory (black dotted line) in phase space starts to deviate from the phase plane diagonal. Black filled circles: stable steady states representing the two choices i.e. choice attractors; grey filled circle: saddle-like unstable steady state. Refer to the main manuscript and previous work[7] for details regarding the content of the phase plane (e.g. nullclines) (**c**) During the middle epoch of the trial, there is a large excitatory feedback from the uncertainty-monitoring monitoring module, such that the phase plane of the sensorimotor module reconfigures, and a new choice-neutral stable steady state appears which aids the initially losing neural population (population 2). The trajectory is now drawn towards this stable steady state, moving back towards the phase plane diagonal. (**d**) During the later epoch of the trial, both sensorimotor populations receive lesser excitatory feedback from the uncertainty-monitoring module, resulting in the phase plane reverting to a similar configuration during the one seen in an early epoch of the trial. It should be noted that a decision is still made by the differential activity amplified by the motor populations, as they continue to integrate excitatory input from the sensorimotor module. Neural population firing rates were calculated by averaging over a time window of 50 ms, slided with a time step of 5 ms.

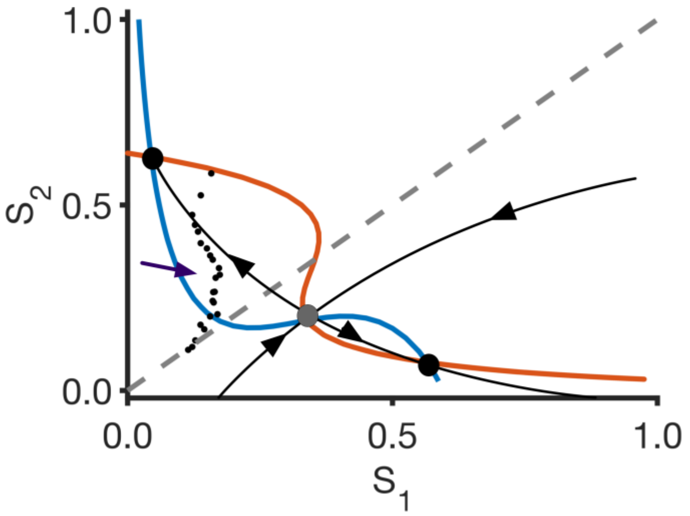


**Supplementary Figure 10. Phase-plane asymmetry of sensorimotor module during an easy task.** Phase space of the sensorimotor module after stimulus onset with evidence quality $\varepsilon = 25.6$. Most of the time, the trajectory will move directly towards the favoured stable steady state i.e. in this case, the attractor representing choice 2/Right. In rare cases where the central choice-neutral stable steady state transiently emerges for such highly biased input, the network would very likely continue its path towards the favoured ('stronger') choice attractor. This is due to the larger basin of attraction of the favoured attractor[7]. The trajectory shown (black dotted line) from stimulus onset to response confirms the dominance of the stable choice attractor albeit the presence of a 'kink' (purple arrow) indicating the transient presence of the central attractor briefly perturbing the trajectory.

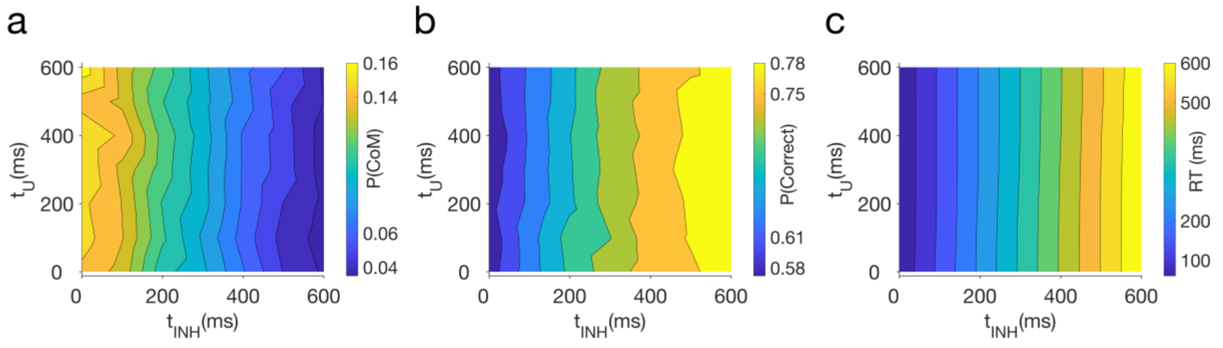


**Supplementary Figure 11. Model performance depends on the time delay in activation onset of the inhibitory neuronal population.** All changes in parameters are simulated with 8000 trials under one condition ($\varepsilon = 6.4\%$). Effects across other difficulty levels can be inferred from the main simulation results (see Fig. 1 and Fig. 4 in manuscript). $t_U$ is the time delay from stimulus onset to activation onset (by removing the top-down inhibition) of the uncertainty-encoding population, while $t_{INH}$ is that for the inhibitory population in the uncertainty-monitoring module. Probability of changes-of-mind P(CoM) decreases, accuracy increases (b) , and response time (RT) is slower (c), as the delay $t_{INH}$ is increased.

| Parameter | Value | Reference, remarks |
|---|---|---|
| $J_{N,ii}$ | 0. 244*nA* | Modified from [7] |
| $J_{N,ij}$ | 0.0497 *nA* | [7] |
| $I_0$ | 0.3255 *nA* | [7] |
| $J_{mc0}$ | 0.009 *nA* $Hz^{-1}$ | Fit to experimental data |
| $J_{A,ext}$ | 0.00052 *nA* $Hz^{-1}$ | [7] |
| $\mu_0$ | 30 Hz | [7] |
| $J_{N,\mathrm{U\_inh}}$ | 0.5 Hz $Hz^{-1}$ | Fit to experimental data |
| $J_{N,LR}$ | 2 *nA* | Modified from [8] |
| $J_{N,RL}$ | 2 *nA* | Modified from [8] |
| $\tau_h$ | *50 ms* | Fit to experimental data |
| $\tau_{\mathrm{mc}}$ | *150 ms* | Fit to experimental data |
| $\tau_{\mathrm{s}}$ | *100 ms* | [7] |
| $a$ | *270 (V nC)*$^{-1}$ | [7] |
| $b$ | *108 Hz* | [7] |
| $d$ | *0.154 s* | [7] |
| $t_{\mathrm{U}}$ | *500 ms* | Fit to experimental data |
| $t_{\mathrm{inh}}$ | *500 ms* | Fit to experimental data |

**Supplementary Table 1: Summary of model parameter values.** See main text for parameter description.
